# Identification of pyroptosis inhibitors that target a reactive cysteine in gasdermin D

**DOI:** 10.1101/365908

**Authors:** Jun Jacob Hu, Xing Liu, Jingxia Zhao, Shiyu Xia, Jianbin Ruan, Xuemei Luo, Justin Kim, Judy Lieberman, Hao Wu

## Abstract

Inflammasomes are multi-protein signalling scaffolds that assemble in response to invasive pathogens and sterile danger signals to activate inflammatory caspases (1/4/5/11), which trigger inflammatory death (pyroptosis) and processing and release of pro-inflammatory cytokines^1,2^. Inflammasome activation contributes to many human diseases, including inflammatory bowel disease, gout, type II diabetes, cardiovascular disease, Alzheimer’s disease, and sepsis, the often fatal response to systemic infection^3–6^. The recent identification of the pore-forming protein gasdermin D (GSDMD) as the final pyroptosis executioner downstream of inflammasome activation presents an attractive drug target for these diseases^7–11^. Here we show that disulfiram, a drug used to treat alcohol addiction^12^, and Bay 11-7082, a previously identified NF-κB inhibitor^13^, potently inhibit GSDMD pore formation in liposomes and inflammasome-mediated pyroptosis and IL-1β secretion in human and mouse cells. Moreover, disulfiram, administered at a clinically well-tolerated dose, inhibits LPS-induced septic death and IL-1β secretion in mice. Both compounds covalently modify a conserved Cys (Cys191 in human and Cys192 in mouse GSDMD) that is critical for pore formation^8,14^. Inflammatory caspases employ Cys active sites, and many previously identified inhibitors of inflammatory mediators, including those against NLRP3 and NF-κB, covalently modify reactive cysteine residues^15^. Since NLRP3 and noncanonical inflammasome activation are amplified by cellular oxidative stress^16–22^, these redox-sensitive reactive cysteine residues may regulate inflammation endogenously, and compounds that covalently modify reactive cysteines may inhibit inflammation by acting at multiple steps. Indeed, both disulfiram and Bay 11-7082 also directly inhibit inflammatory caspases and pleiotropically suppress multiple processes in inflammation triggered by both canonical and noncanonical inflammasomes, including priming, puncta formation and caspase activation. Hence, cysteine-reactive compounds, despite their lack of specificity, may be attractive agents for reducing inflammation.

---

We performed high-throughput screening to discover inhibitors of GSDMD using a fluorogenic liposome leakage assay, which detects leakage of Tb^3+^ from Tb^3+^-loaded liposomes incubated with GSDMD and caspase-11^7–9^ (Fig. 1a). Concentrations of liposomes, caspase-11 and GSDMD were optimized to achieve a Z’ value of ~0.7, a cutoff that provides reproducible separation of hits from controls^23^ (Extended Data Fig. 1a-c). We screened 3,752 small molecules from a Harvard ICCB-Longwood collection to look for compounds that inhibited liposome leakage by at least 50% (Fig. 1b). After excluding pan-assay-interference compounds that non-specifically react with many biological targets and GSDMD-independent quenchers of fluorescence, we identified 22 active compounds and measured their IC_50_ values. The most potent inhibitor was C-23, which had an IC_50_ of 0.30 ± 0.01 µM (Fig. 1c-e, Extended Data Fig. 1d). C-23 is a symmetrical molecule known as disulfiram, a drug used to treat alcohol addiction^12^. C-22, -23 and -24 were selected for further studies based on their low IC_50_ values and GSDMD binding, assessed by microscale thermophoresis (MST) (Fig. 1c,f).

**Figure 1.**
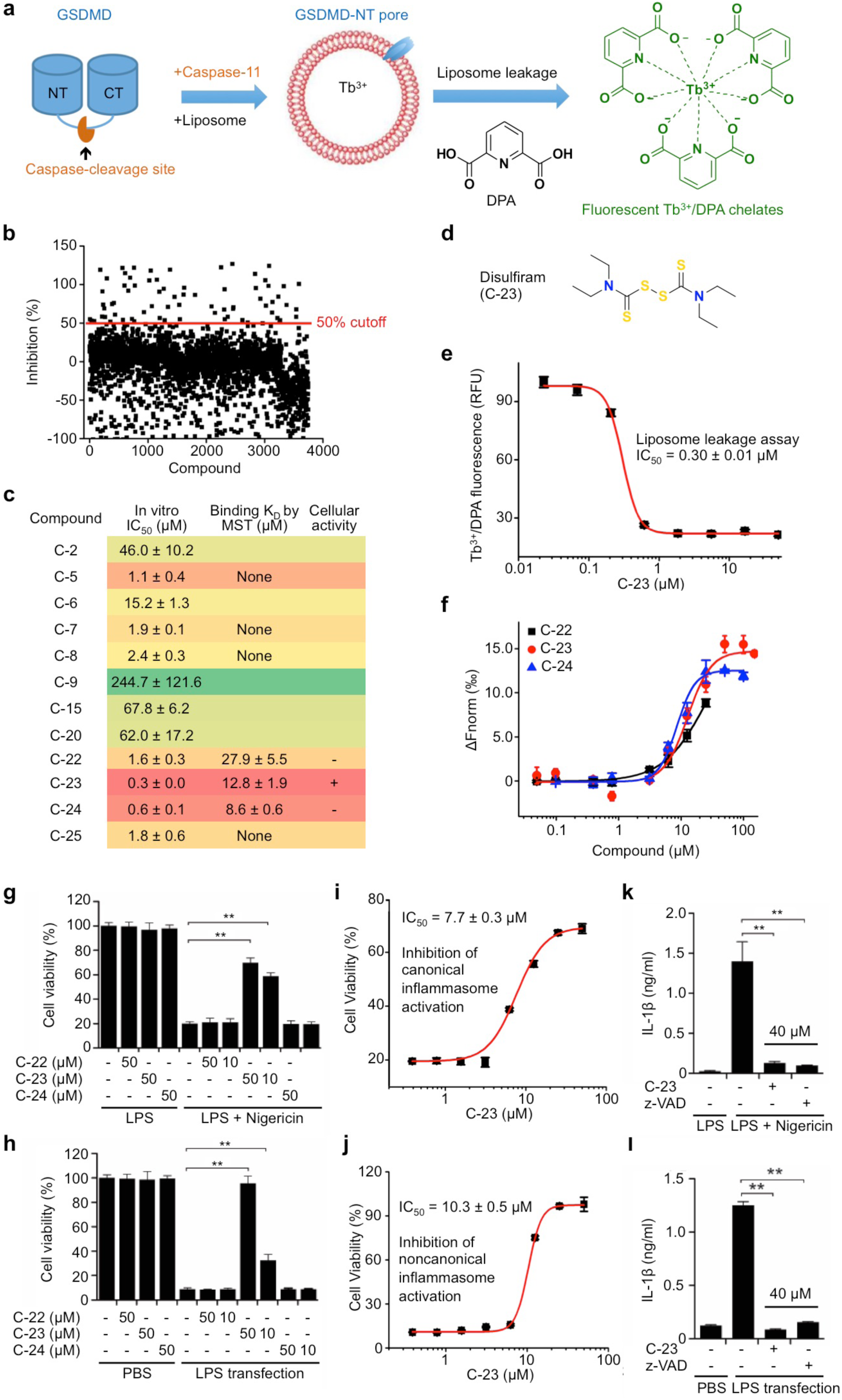
High throughput screen identifies disulfiram as an inhibitor of GSDMD pore formation. **a**, Pictorial representation of the terbium (Tb^3+^)/dipicolinic acid (DPA) fluorescence liposome leakage assay. **b**, Percentage inhibition of liposome leakage by each compound, assayed at 25 μg/mL (~50 μM for most compounds). Cutoff was 50% inhibition. **c**, IC50 values of the 12 screening hits after excluding Tb3+/DPA assay quenchers and hits without saturable IC_50_ curve. The top 7 hits were assessed for GSDMD binding by microscale thermophoresis (MST) (see **f**) and compounds that bound were tested for pyroptosis inhibition in cells (see **g,h**). **d**, Chemical structure of compound C-23. **e**, Dose response curve of compound C-23 in liposome leakage assay. **f**, MST measurement of the binding of Alexa 488-labeled His-MBP-GSDMD (80 nM) with C-22, C-23 or C-24. **g**,**i**,**k**, PMA-differentiated LPS-primed human THP-1 were pretreated with indicated concentrations of each compound for 1 h before adding nigericin or medium. The number of surviving cells was determined by CellTiter-Glo assay (**g,i**) and IL-1b in culture supernatants was assessed by ELISA (**k**) 2 hrs later. **h**,**j**,**l**, Mouse iBMDMs were pretreated with each compound for 1 hr before electroporation with PBS or LPS. The number of surviving cells was determined by CellTiter-Glo assay **(h,j)** and IL-1b in culture supernatants was assessed by ELISA (**l)** 2.5 hrs later. In **(k,l)** 40 μM inhibitors were added. Graphs show the mean ± s.d. and data shown are representative of three independent experiments. **P < 0.01.

To evaluate whether C-22, -23, and -24 inhibit pyroptosis, we added these compounds to PMA-differentiated and LPS-primed human THP-1 cells or mouse immortalized bone marrow-derived macrophages (iBMDMs) before activating the canonical inflammasome with nigericin or the non-canonical inflammasome by LPS electroporation. Only C-23 (disulfiram) blocked pyroptosis in cells, with IC_50_ values of 7.67 ± 0.29 µM and 10.33 ± 0.50 µM for canonical and non-canonical inflammasome-dependent pyroptosis, respectively (Fig. 1g-j), and impaired cell death triggered by the AIM2 inflammasome in mouse iBMDMs transfected with poly(dA:dT) (Extended Data Fig. 1e). Disulfiram also inhibited nigericin-or LPS transfection-induced IL-1β secretion with potency comparable to the pan-caspase inhibitor z-VAD-fmk (Fig. 1k,l).

Disulfiram is being investigated as an anticancer agent because epidemiological studies showed that individuals taking disulfiram for alcohol addiction were less likely to die of cancer^24^. In cells disulfiram is rapidly metabolized to diethyldithiocarbamate (DTC)^25,26^. The anti-cancer activity of DTC in vivo is greatly enhanced by complexation with copper^24^, likely because of the enhanced electrophilicity of the DTC thiols. In liposome leakage assay, we found that copper gluconate (Cu^2+^) only weakly increased disulfiram or DTC inhibition (Fig. 2a); we interpret this as due to the high reactivity of the GSDMD Cys residue involved (see below). However, Cu^2+^ strongly promoted the ability of either disulfiram or DTC to protect LPS-primed THP-1 cells from pyroptosis (Fig. 2b). With Cu^2+^, the IC_50_ of disulfiram for inhibiting pyroptosis decreased 24-fold to 0.41 ± 0.02 µM, which was similar to its potency for preventing liposome leakage. DTC became almost as active as disulfiram in cells in the presence of Cu^2+^.

**Figure 2.**
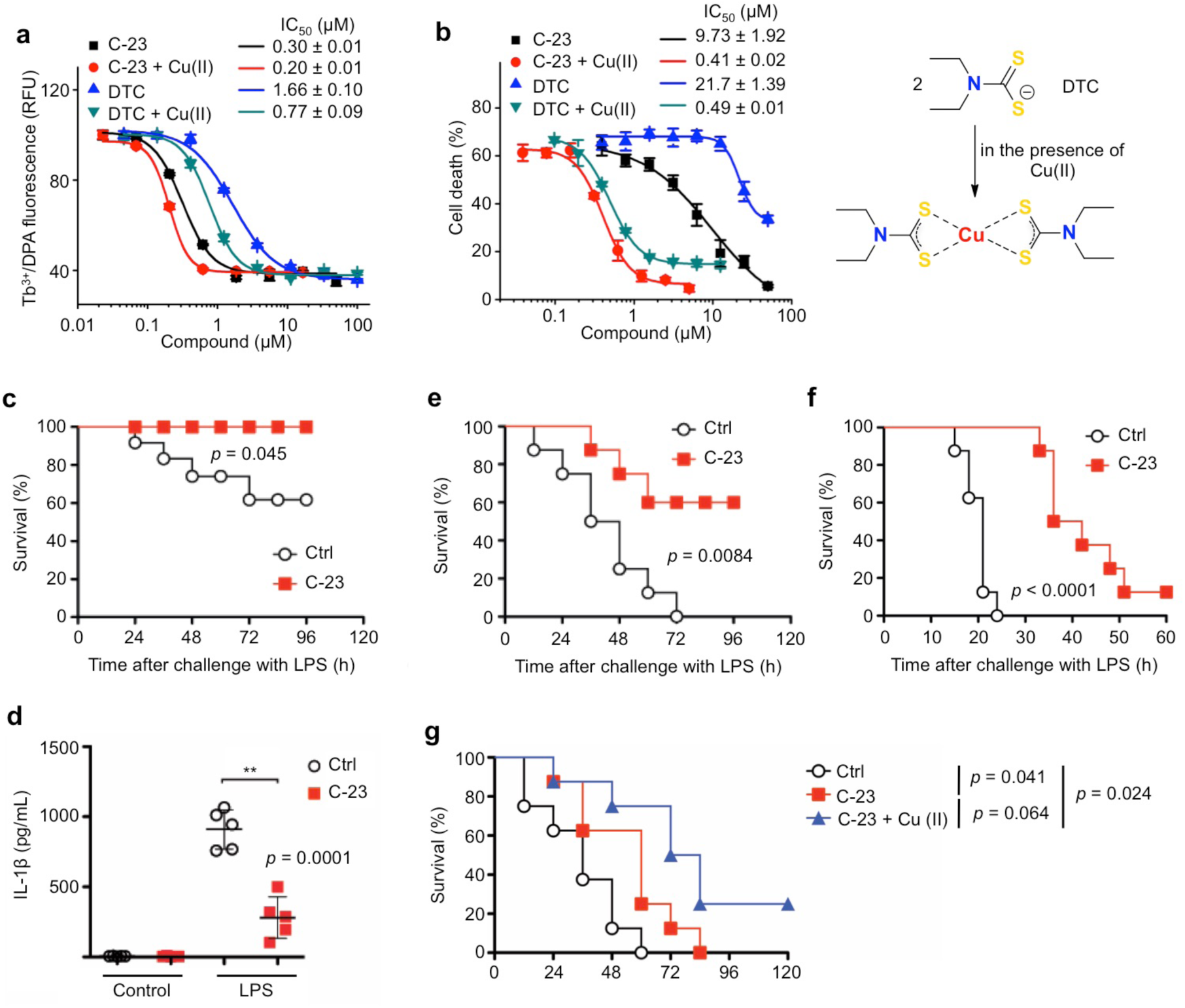
Disulfiram activity in cells is enhanced by Cu^2+^ and disulfiram protects against LPS-induced sepsis. **a**, Dose response curves of inhibition of liposome leakage by disulfiram (C-23) or its metabolite DTC in the presence or absence of Cu(II). **b**, LPS-primed THP-1 were pretreated with C-23 or DTC in the presence or absence of Cu(II) for 1 hr before adding nigericin or medium for 2 hrs. Cell death was determined by CytoTox96 assay. **c,d,e,f,** Mice were pretreated with C-23 (50 mg/kg) or vehicle (Ctrl) by intraperitoneal injection 24 and 4 hrs before intraperitoneal LPS challenge (**c**,**d,** 15 mg/kg; **e**, 25 mg/kg; **f**, 50 mg/kg) and followed for survival. Statistical analysis was performed using the log-rank test (**c,e,f,** 8 mice/group). **d**, Serum IL-1b measured by ELISA in mice (n = 5/group) pretreated with disulfiram as above and challenged with 15 mg/kg LPS. Serum was obtained 12 hrs post LPS challenge. Shown are mean ± s.d. **g**, Mice were treated with C-23 (50 mg/kg), C-23 (50 mg/kg) plus copper gluconate (0.15 mg/kg) or vehicle (Ctrl) by intraperitoneal injection 0 and 12 hrs post intraperitoneal LPS challenge (25 mg/kg). Statistical analysis was performed using the log-rank test (8 mice/group).

Because disulfiram inhibited pyroptosis and IL-1β release in cells, we next tested its ability to protect C57BL/6 mice from LPS-induced sepsis. Mice were treated with vehicle or disulfiram intraperitoneally before challenge with LPS. Whereas the lowest concentration of LPS (15 mg/kg) killed 3 of 8 control mice after 96 hrs, all the disulfiram-treated mice survived (P < 0.05) (Fig. 2c). Serum IL-1β concentrations were strongly reduced 12 hrs after LPS challenge when all mice were alive (281 ± 149 ng/mL in disulfiram-pretreated mice, 910 ± 140 ng/mL in control mice (P < 0.0001)) (Fig. 2d). Following LPS challenge at the intermediate concentration (25 mg/kg), all the control mice died within 72 hrs, but 5 of 8 of the disulfiram-treated mice survived (P < 0.01) (Fig. 2e). At the highest LPS challenge (50 mg/kg), while all the control mice died within a day, death was significantly delayed by disulfiram treatment and 1 of 8 mice survived (P < 0.0001) (Fig. 2f). To determine if we could delay treatment until after LPS challenge and whether adding copper could improve protection, we challenged mice with 25 mg/kg LPS intraperitoneally and administered disulfiram with or without copper gluconate immediately and 24 hrs later. Post-LPS treatment still improved survival (P = 0.041 without copper and P = 0.024 with copper). All the control mice and mice treated without copper died, but 2 of 8 mice given copper-complexed disulfiram survived (Fig. 2g). Thus, disulfiram given before or after LPS partially protected mice from septic death and reduced IL-1β secretion.

Disulfiram has been shown to inactivate reactive Cys residues by covalent modification^27^. To probe the mechanism of GSDMD inhibition by disulfiram, we used nano-liquid chromatography-tandem mass spectrometry (nano-LC-MS/MS) to analyse disulfiram-treated human GSDMD. Tryptic fragments indicated a dithiodiethylcarbamoyl adduct of Cys191, in which half of the symmetrical disulfiram molecule is attached to the thiol (Fig. 3a,b, Extended Data Fig. 2). Indeed, Cys191 is required for GSDMD pore formation in cells, since oligomerization was blocked by Ala mutation of the corresponding Cys192 in mouse GSDMD^8^. This Cys residue, conserved in GSDMD, but not in other GSDM family members, is accessible in both the full-length autoinhibited structure model and the N-terminal pore form model, generated based on mouse GSDMA3 structures^7,14^ (Fig. 3c, Extended Data Fig. 3a). Corresponding to Leu183 of GSDMA3, Cys191 sits at the distal tip of the membrane spanning region at the beginning of the β8 strand within the β7-β8 hairpin, which is a key element in the β-barrel that forms the pore^14^. Analysis of Cys reactivity using PROPKA^28^ suggests that Cys191 is the most reactive among all Cys residues in GSDMD. Consistent with its high reactivity, a time course analysis showed that disulfiram inhibited liposome leakage within 2 min of incubation (Extended Data Fig. 3b). To confirm that disulfiram acts on Cys191, we generated Ala mutations of Cys191, and of Cys38 as a control. Whereas the disulfiram IC_50_ values for WT and C38A were both around 0.3 µM in the liposome leakage assay, the IC_50_ for C191A was about 8-fold higher (Fig. 3d). We also incubated disulfiram with N-acetylcysteine (NAC), which contains a reactive Cys that can inactivate Cys-reactive drugs, before assessing whether disulfiram protects THP-1 cells from nigericin-mediated pyroptosis. As expected, NAC eliminated the activity of disulfiram (Fig. 3e). These data together suggest that disulfiram inhibits GSDMD pore formation by selectively and covalently modifying Cys191.

**Figure 3.**
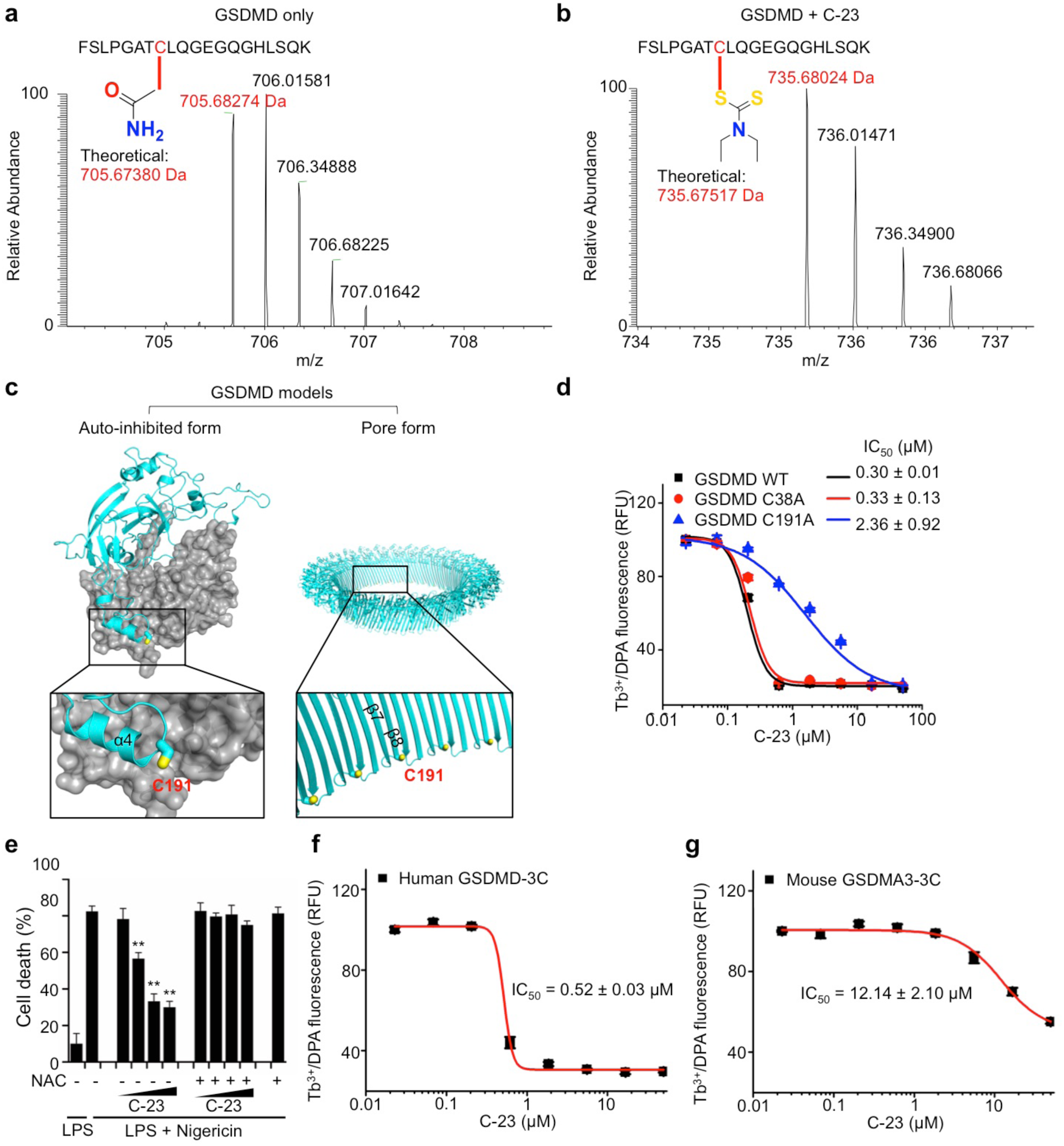
Disulfiram covalently modifies GSDMD Cys191. **a**,**b**, MS/MS spectra of the Cys191-containing human GSDMD peptide FSLPGAT**C**LQGEGQGHLSQK (aa 184-103; 2057.00 Da) modified on Cys191 (red) by carbamidomethyl (an increase of 57.0214 Da) [LC retention time, 22.85 min; a triplet charged precursor ion m/z 705.6827 (mass: 2114.0481 Da; delta M 2.27 ppm) was observed] (**a**) or of the corresponding GSDMD peptide after GSDMD incubation with C-23 (disulfiram), which was modified on Cys191 (red) by the diethyldithiocarbamate moiety of C-23 (an increase of 147.0255 Da). [LC retention time. 28.93 min; a triplet charged precursor ion m/z 735.6802 (mass: 2204.0406 Da; delta M 0.53 ppm) was observed.] (**b**). **c**, Models of full-length human GSDMD in its auto-inhibited form and of the pore form of GSDMD N-terminal fragment (GSDMD-NT) based on the corresponding structures of GSDMA3 ^7,14^ showing the location in yellow of Cys191, modified by compound C-23. GSDMD-NT in cyan; GSDMD-CT in grey. **d**, Dose response curve of C-23 inhibition of liposome leakage induced by wild-type, C38A or C191A GSDMD (0.3 μM) plus caspase-11 (0.15 μM). **e**, C-23 inhibition of pyroptosis of LPS + nigericin treated THP-1 cells after C-23 preincubation for 1 hr with N-acetylcysteine (NAC, 500 μM) or medium. 2-fold dilutions of C-23 ranging from 5 to 40 μM were used. Graphs show the mean ± s.d. and data shown are representative of three independent experiments. **P < 0.01. **f**,**g**, Dose response curve of compound C-23 in liposome leakage induced by human GSDMD-3C (0.3 μM) plus 3C protease (0.15 μM) (**f**) or mouse GSDMA3-3C (0.3 μM) plus 3C protease (0.15 μM) (**g**).

Disulfiram has been reported to inhibit caspases by binding to the catalytic Cys responsible for proteolysis^29^. It is therefore likely that disulfiram inhibits both caspases and GSDMD. Using a fluorogenic caspase activity assay that measures the release of 7-amino-4-methylcoumarin (AMC) from substrate Ac-YVAD-AMC, we found that disulfiram indeed inhibited caspase-1 and caspase-11 (Extended Data Fig. 4). To determine the relative contribution of caspase-11 inhibition versus GSDMD inhibition by disulfiram in pore formation, we replaced the caspase cleavage site in GSDMD with the rhinovirus 3C protease site (GSDMD-3C) and used the 3C protease instead of caspase-11 in the liposome leakage assay. The resulting IC_50_ was 0.52 ± 0.03 µM, comparable to 0.30 ± 0.01 µM for caspase-11-triggered liposome leakage (Fig. 1e, Fig. 3f). By contrast, as mouse GSDMA3 lacks the conserved Cys191, disulfiram inhibited liposome leakage triggered by 3C-cleaved GSDMA3 containing a 3C protease site (GSDMA3-3C) with a much weaker IC_50_ of 12.14 ± 2.10 µM (Fig. 3g). Thus, we conclude that the inhibitory effect of disulfiram in the liposome leakage assay is mainly mediated by direct inhibition of GSDMD.

**Figure 4.**
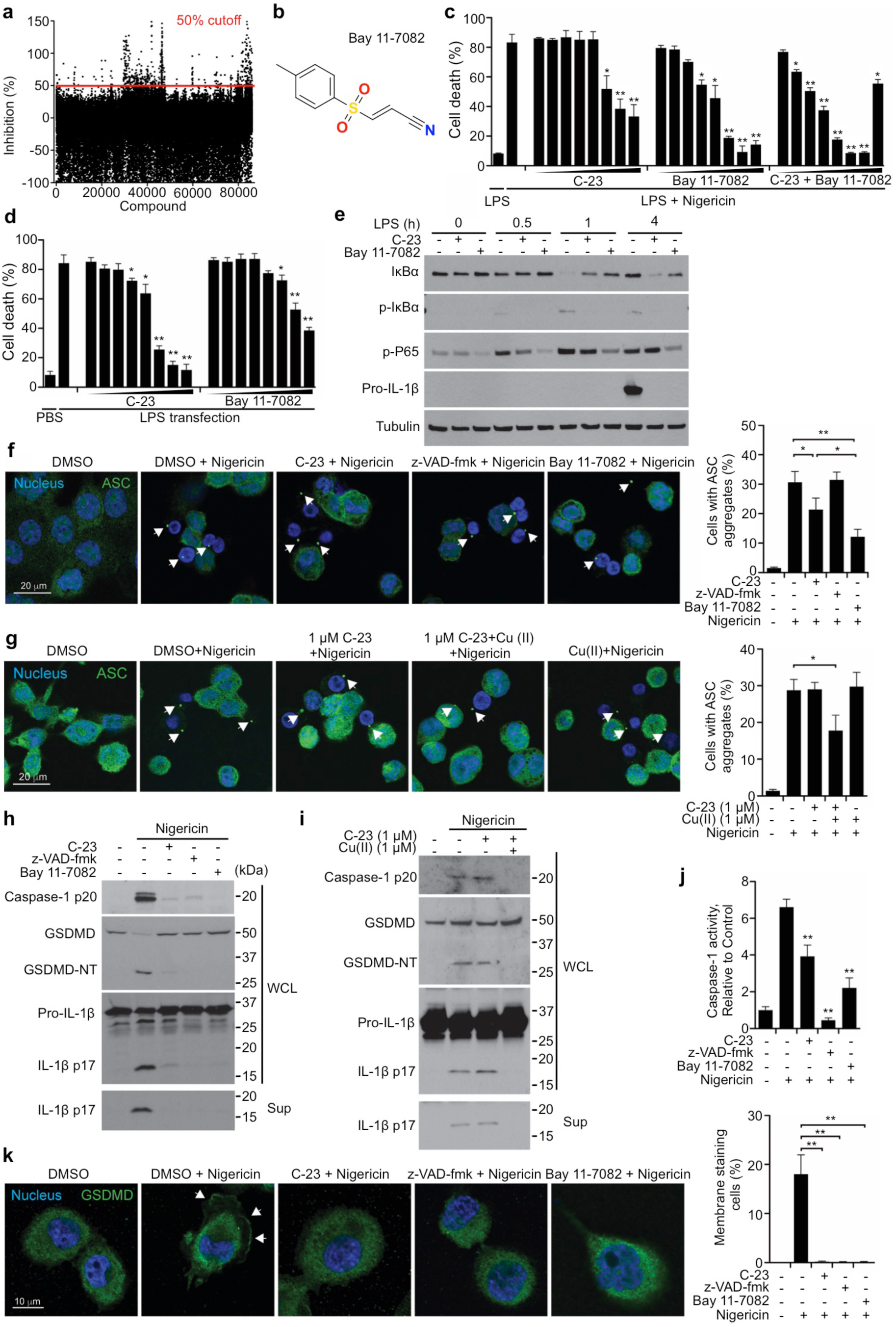
Disulfiram and Bay 11-7082 inhibit multiple steps in inflammasome activation. **a**, Percentage inhibition of liposome leakage for each of the 86,050 compounds assayed in the second screen at 25 μg/mL (~50 μM for most compounds). Cutoff was 50% inhibition. **b**, Chemical structure of compound Bay 11-7082. **c**, PMA-differentiated LPS-primed THP-1 cells were pretreated with 2-fold serial dilutions (ranging from 0.3125 to 40 μM) of C-23 and/or Bay 11-7082 for 1 hr before treatment with nigericin. Cell death was determined by CytoTox96 assay. **d**, Mouse iBMDMs were pretreated with serial 2-fold dilutions of C-23 or Bay 11-7082 (ranging from 0.3125 to 40 μM) for 1 hr before electroporation with PBS or LPS. Cell death was determined by CytoTox96 assay. **e**, THP-1 cells were pretreated with 30 μM C-23 or Bay 11-7082 for 1 hr before adding LPS. Shown are immunoblots of whole cell lysates harvested 0.5 hr later. **f**,**h**,**j**, LPS-primed THP-1 were pretreated with 30 μM C-23, Bay 11-7082 or z-VAD-fmk for 1 hr before adding nigericin or medium. Representative images of ASC specks (arrowheads) (left) and mean ± s.d. percent of cells with ASC specks (right) analysed 20 min later (**f**). Whole cell lysates (WCL) and culture supernatants (Sup) were harvested 30 min after adding nigericin and immunoblotted with the indicated antibodies (**h**). Caspase-1 activity was assayed 30 min after adding nigericin using a cell-permeable fluorescence dye FAM-YVAD-FMK (**j**). **g,i,** LPS-primed THP-1 were pretreated with 1 μM C-23 in the presence or absence of Cu(II) for 1 hr before adding nigericin or medium. Representative images of ASC specks (arrowheads) (left) and mean ± s.d. percent of cells with ASC specks (right) analysed 20 min later (**g**). Whole cell lysates (WCL) and culture supernatants (Sup), harvested 30 min after adding nigericin, were analyzed by immunoblot (**i**). **k**, LPS-primed THP-1 were pretreated with 30 μM C-23, Bay 11-7082 or z-VAD-fmk for 1 hr before adding nigericin or medium and stained with a mouse anti-GSDMD monoclonal antibody (see Extended Data Fig. 6) 30 min later. Representative confocal microscopy images (left) and quantification (right) of proportion of cells with GSDMD membrane staining and pyroptotic bubbles. Arrows indicate GSDMD staining of pyroptotic bubbles. Graphs show the mean ± s.d; data are representative of three independent experiments. *P < 0.05, **P < 0.01.

To determine the structure-activity relationship (SAR) of disulfiram, we evaluated a set of disulfiram analogues and found that a number of small alkyl-substituted thiuram disulfides had IC_50_ values in liposome leakage assay comparable to or marginally better than disulfiram (Extended Data Fig. 5a,b). Potency was mildly negatively correlated with the steric bulkiness of the substituents, presumably due to the need for the core thiuram disulfide chemical motif to access the reactive Cys (Extended Data Fig. 5a,b). Several small alkyl-substituted thiuram disulfides also significantly protected against nigericin-induced pyroptosis in THP-1 cells, albeit less potent than disulfiram (Extended Data Fig. 5c-e).

To identify additional inhibitors of pyroptosis, we expanded our liposome leakage screen to test 86,050 additional compounds in the ICCB-Longwood collection. 343 hit compounds inhibited liposome leakage by at least 50% (Fig. 4a). However, when these were tested in a high throughput cell viability assay for inhibition of the canonical inflammasome pathway in THP-1 cells, only 2 compounds inhibited cell death by ≥ 50%. One was the pan-caspase inhibitor z-VAD-fmk and the other was Bay 11-7082, a previously known inhibitor of NF-κB activation^13^ and the NLRP3 pathway^30^ (Fig. 4b,c). Bay 11-7082 bound to GSDMD according to MST, but was less active at inhibiting liposome leakage than disulfiram (IC_50_ 6.81 ± 0.10 µM vs 0.30 ± 0.01 µM) (Extended Data Fig. 6a, b and Fig. 1e). Bay 11-7082 inhibited caspase-1 similarly to disulfiram, but was about 3 times less active against caspase-11 (Extended Data Fig. 4a-d, Extended Data Fig. 6a-d). Surprisingly, like disulfiram, Bay 11-7082 functions by inactivating reactive Cys residues^31,32^, and Cys191 in GSDMD was covalently modified by Bay 11-7082 (Extended Data Fig. 6e,f). However, Bay 11-7082 inhibition of liposome leakage was only reduced 2-fold by substituting C191A GSDMD for WT GSDMD in the liposome leakage assay (Extended Data Fig. 6a). Indeed, much of Bay 11-7082 inhibition of liposome leakage could be attributed to caspase-11 inhibition, since Bay 11-7082 was substantially less able to inhibit leakage caused by GSDMD-3C plus 3C protease than by GSDMD plus caspase-11 and its activity against mouse GSDMA3-3C, which lacks a comparable reactive cysteine, plus 3C protease was similar to its activity against GSDMD-3C (Extended Data Fig. 6g,h).

Bay 11-7082 inhibited pyroptosis triggered by both the canonical and non-canonical inflammasomes in THP-1 cells, but was more active in nigericin-treated than LPS-transfected cells (Fig. 4c, d). Bay 11-7082 was more effective at inhibiting canonical inflammasome-dependent pyroptosis than disulfiram in the absence of copper, and the two drugs together had an additive protective effect, although were cytotoxic at the highest concentration tested (Fig. 4c). However, Bay 11-7082 was less active than disulfiram at inhibiting pyroptosis induced by non-canonical inflammasome activation (Fig. 4d).

Because both disulfiram and Bay 11-7082 non-specifically modify reactive Cys, we next analysed their effects on the steps leading to pyroptosis and inflammatory caspase activation. Some of the genes that participate in the canonical inflammasome pathway are not expressed in unstimulated cells and their expression needs to be induced, often by binding to cell surface sensors of pathogen and danger-associated molecular patterns, such as Toll-like receptors (TLR), in a process called priming. Bay 11-7082 is known to inhibit NF-κB activation, a key transcription factor in priming. We first looked at the effect of disulfiram and Bay 11-7082 on priming (Fig. 4e). NF-κB activation was assessed by examining IκBα phosphorylation and degradation and RelA (p65) phosphorylation. Induction of pro-IL-1β was assessed by immunoblot for pro-IL1β protein. In the absence of disulfiram or Bay 11-7082, phosphorylation of p65 was first detected 30 min after adding LPS and persisted for 4 hrs, phosphorylation and reduced IκBα were detected 1 hr after adding LPS, and increased pro-IL-1β was detected 4 hrs after adding LPS. Both drugs, added at 30 µM concentrations, inhibited NF-κB activation, but Bay 11-7082 had a stronger effect; both blocked pro-IL-1β induction. Thus, disulfiram and Bay 11-7082 both inhibit priming.

Nigericin activates the assembly of the NLRP3 canonical inflammasome using an adaptor called apoptosis-associated speck-like protein containing a caspase recruitment domain (ASC), which can be visualized in immunofluorescent microscopy as specks. When LPS-primed THP-1 cells were treated with nigericin in the absence of inhibitors, ASC specks were detected in 30% of cells (Fig. 4f). As expected, speck formation was not inhibited by z-VAD-fmk, since caspase activation occurs downstream of inflammasome assembly. However, both drugs, added after priming but one hour before nigericin, inhibited ASC speck formation, but not completely, and Bay 11-7082 was more potent than disulfiram when used at the same concentration. 1 µM disulfiram was completely inactive at blocking pyroptosis triggered by nigericin or transfected LPS (Fig. 1i,j), but the same concentration of disulfiram in combination with copper gluconate blocked pyroptosis completely and also reduced ASC puncta (Fig. 4g).

Canonical inflammasome activation activates caspase-1, which cleaves pro-IL-1β and GSDMD, which forms pores needed to release processed IL-1β. To assess which steps in NLRP3-mediated inflammation were inhibited post ASC speck formation, LPS-primed THP-1 cells were treated with vehicle or 30 µM z-VAD-fmk, disulfiram or Bay 11-7082 1 hr before adding nigericin, and cleavage and activation of caspase-1, GSDMD, and pro-IL-1β were analysed by immunoblot of whole cell lysates 30 min later (Fig. 4h). Secretion of processed IL-1β was also assessed by immunoblot of culture supernatants. Caspase-1, GSDMD and pro-IL-1β cleavage to their active forms was clearly detected in the absence of inhibitors, but was dramatically reduced in cells treated by any of the 3 inhibitors; moreover, processed IL-1β was only detected in the culture supernatants in the absence of any inhibitor. When the same experiment was repeated by treating cells with only 1 µM disulfiram in PBS or copper gluconate, disulfiram complexed with copper completely blocked caspase-1, GSDMD, and pro-IL-1β processing and IL-1β secretion, but disulfiram without copper had no effect (Fig. 4i). Because immunoblots are not quantitative, we also assessed caspase-1 activity 30 min after adding nigericin using a fluorescent substrate in intact cells. While caspase-1 activity was completely inhibited by z-VAD-fmk, it was only partially reduced by either disulfiram and Bay 11-7082, again more strongly by Bay 11-7082 (Fig. 4j). Next we assessed the effect of z-VAD-fmk, disulfiram and Bay 11-7082 on LPS + nigericin-induced GSDMD pore formation by immunofluorescence microscopy using a monoclonal antibody we generated that recognizes both uncleaved GSDMD and its pore form (Fig. 4k, Extended Data Fig. 7). In the absence of any inhibitor, the GSDMD antibody stained both the cytosol and the plasma membrane of LPS plus nigericin treated cells, which formed characteristic pyroptotic bubbles^10^. All 3 inhibitors completely blocked GSDMD membrane staining and the appearance of pyroptotic bubbles. Thus, disulfiram and Bay 11-7082 inhibit multiple steps leading to canonical inflammasome-induced pyroptosis and inflammatory cytokine release, including priming, inflammasome assembly, inflammatory caspase activation, pro-inflammatory cytokine processing and GSDMD pore formation.

Our identification of Cys191-modifying compounds that inhibit pore formation suggests that Cys191 is also likely modified in cells by endogenous agents, which may regulate pore formation in a redox-sensitive manner. Cysteine-reactive drugs are notoriously non-specific as to their targets. Indeed, disulfiram is used to treat chronic alcohol addiction, where it inhibits acetaldehyde dehydrogenase^12^, and is also known to inhibit multiple protein phosphatases and caspases^29,33^. Nonetheless, disulfiram is a very safe drug even at high concentrations and has a very long half-life in tissues. We speculate that the promiscuity of disulfiram allows it to react preferentially with different targets under different conditions, e.g. with acetaldehyde dehydrogenase in alcohol addiction, or with inflammasome components, including GSDMD, during inflammation. Perhaps its lack of toxicity may be because of incomplete inhibition of individual targets that tune down a response without causing side effects. In inflammation its effectiveness may be amplified by simultaneously disrupting multiple inflammatory steps. Indeed, in our experiments, inhibition of upstream steps in inflammation, such as ASC puncta formation, tended to be partial, while inhibition of the ultimate cellular events, pyroptosis and inflammatory cytokine secretion, were more dramatic.

Inflammation appears to be especially sensitive to Cys-modifying drugs. Of note, many of the compounds that have been previously identified to inhibit other steps in priming and activating inflammasomes work by covalently binding to reactive Cys residues in inflammation pathway molecules, including the inflammatory caspase inhibitors, the diarylsulfonylureas that inhibit inflammatory cytokine release, MCC950 and 3,4-methylenedioxy-β-nitrostyrene that inactivate NLRP3, and myochrysine and related compounds that inhibit IKKβ^15^. The repeated identification of cysteine-modifying drugs as inhibitors of inflammation suggests that many inflammatory pathway molecules have reactive Cys residues, which are highly sensitive to oxidation or modification, such as Cys191 of human GSDMD that we identified here. These results in their ensemble raise the possibility that inflammation pathways are regulated by the oxidative state of a cell, namely by the generation of reactive oxygen species, and by cellular levels of antioxidants like glutathione and thioredoxin. In fact, oxidized endogenous lipids are known to regulate the non-canonical inflammasome^19–21^. A number of papers have implicated oxidation in the most studied inflammatory pathway, mediated by the NLRP3 inflammasome^16–18,22^. However, the role of oxidation in inflammasome activation is controversial, suggesting that the overall role of oxidants and antioxidants may be a complex summation of positive and negative feedback effects that may vary in different cell types or with different inflammatory instigators. One wonders how oxidative events that are triggered by infection or by the oxidative burst in response to infection in myeloid cells impact the extent of inflammation. The role of oxidation and endogenous and exogenous anti-oxidants in inflammation, suggested by the repeated identification of critical reactive Cys in key inflammatory mediators, merits further study.

## Methods

### Mice

8-week-old female C57BL/6 wild-type mice were purchased from The Jackson Laboratory and maintained at the SPF facility at Harvard Medical School. All mouse experiments were conducted using protocols approved by the Animal Care and Use Committees of Boston Children’s Hospital and Harvard Medical School.

### Reagents

β-mercaptoethanol (2ME), dithiothreitol (DTT), terbium(III) chloride (TbCl_3_), dipicolinic acid (DPA) and copper gluconate were from Sigma-Aldrich. Compound C-23 and its analogues: Tetraethylthiuram disulfide (C-23), tetramethylthiuram disulfide (C-23A1), tetrabutylthiuram disulfide (C-23A3), 4-Methylpiperazine-1-carbothioic dithioperoxyanhydride (C-23A4), Tetraphenylthiuram disulfide (C-23A5), *N*,*N*’-Dimethyl-*N*,*N*’-(4,4’-dimethyldiphenyl)thiuram disulfide (C-23A6), di(4-morpholinyl)dithioperoxyanhydride (C-23A7), *N*,*N*’-Dimethyl-*N*,*N*’-di(4-pyridinyl)thiuram disulfide (C-23A8), pyrrolidine-1-carbothioic dithioperoxyanhydride (C-23A10), and dimethyldiphenylthiuram disulfide (C-23A11) were from Sigma-Aldrich. Tetraisopropylthiuram disulfide (C-23A2) and dicyclopentamethylenethiuram disulfide (C-23A9) were from Oakwood Chemicals. Tetrabenzylthiuram disulfide (C-23A12) was from AK Scientific. Phorbol 12-myristate 13-acetate (PMA) and DMSO were from Sigma-Aldrich. Ultra LPS and nigericin were from InvivoGen. The pan-caspase inhibitor z-VAD-fmk was from BD Bioscience. The complete protease inhibitor cocktail and the PhosSTOP phosphatase inhibitor cocktail were from Roche.

The monoclonal antibody against GSDMD was generated in house by immunizing 6 week-old BALB/c mice with recombinant human GSDMD and boosting with recombinant human GSDMD-NT according to standard protocols. Serum samples were collected to assess titers of reactive antibodies and spleen cells were fused with SP2/0 myeloma cells. Hybridomas were selected and supernatants from the resulting clones were screened by enzyme linked immunosorbent assay (ELISA), immunoblot and immunofluorescence microscopy. Tubulin antibody was from Sigma-Aldrich. Phospho-IκBα antibody, IκBα antibody, Phospho-NF-κB p65 antibody, and cleaved human caspase-1 (Asp297) antibody were from Cell Signaling Technology. ASC antibody (AL177) and mouse caspase-1 p20 antibody were from AdipoGen. Human and mouse IL-1β antibodies were from R&D Systems.

### Protein expression and purification

Full-length human GSDMD sequence was cloned into the pDB.His.MBP vector with a tobacco etch virus (TEV)-cleavable N-terminal His_6_-MBP tag using NdeI and XhoI restriction sites. Human GSDMD-3C and mouse GSDMA3-3C mutants were constructed by QuikChange Mutagenesis (Agilent Technologies). For expression of full-length GSDMD, GSDMD-3C, GSDMA3, and GSDMA3-3C, *E. coli* BL21 (DE3) cells harbouring the indicated plasmids were grown at 18 °C overnight in LB medium supplemented with 50 µg ml^−1^ kanamycin after induction with 0.5 mM isopropyl-β-D-thiogalactopyranoside (IPTG) when OD_600_ reached 0.8. Cells were ultrasonicated in lysis buffer containing 25 mM Tris-HCl at pH 8.0, 150 mM NaCl, 20 mM imidazole and 5 mM 2ME. The lysate was clarified by centrifugation at 40,000xg at 4 °C for 1 hr. The supernatant containing the target protein was incubated with Ni-NTA resin (Qiagen) for 30 min at 4 °C. After incubation, the resin–supernatant mixture was poured into a column and the resin was washed with lysis buffer. The protein was eluted using the lysis buffer supplemented with 100 mM imidazole. The His_6_-MBP tag was removed by overnight TEV protease digestion at 16 °C. The cleaved protein was purified using HiTrap Q ion-exchange and Superdex 200 gel-filtration columns (GE Healthcare Life Sciences).

Caspase-11 sequence was cloned into the pFastBac-HTa vector with a TEV cleavable N-terminal His_6_-tag using EcoRI and XhoI restriction sites. The baculoviruses were prepared using the Bac-to-Bac system (Invitrogen), and the protein was expressed in Sf9 cells following the manufacturer’s instructions. His–caspase-11 baculovirus (10 ml) was used to infect 1 L of Sf9 cells. Cells were collected 48 hrs after infection and His_6_–caspase-11 was purified following the same protocol as for His_6_-MBP–GSDMD. Eluate from Ni-NTA resin was collected for subsequent assays.

### Liposome preparation

PC (1-palmitoyl-2-oleoyl-sn-glycero-3-phosphocholine, 25 mg/mL in chloroform; 80 µL), PE (1-palmitoyl-2-oleoyl-sn-glycero-3-phosphoethanolamine, 25 mg/mL in chloroform; 128 µL) and CL (1’,3’-bis[1,2-dioleoyl-*sn*-glycero-3-phospho]-*sn*-glycerol (sodium salt), 25 mg/mL in chloroform; 64 µL) were mixed and the solvent was evaporated under a stream of N_2_ gas. The lipid mixture was suspended in 1 mL Buffer A (20 mM HEPES, 150 mM NaCl, 50 mM sodium citrate, and 15 mM TbCl_3_) for 3 min. The suspension was pushed through 100 nm Whatman^®^ Nuclepore™ Track-Etched Membrane 30 times to obtain homogeneous liposomes. The filtered suspension was purified by size exclusion column (Superose 6, 10/300 GL) in Buffer B (20 mM HEPES, 150 mM NaCl) to remove TbCl_3_ outside liposomes. Void fractions were pooled to produce a stock of PC/PE/CL liposomes (1.6 mM). The liposomes are diluted to 50 µM with Buffer C (20 mM HEPES, 150 mM NaCl and 50 µM DPA) for use in high-throughput screening.

### High-throughput screen for GSDMD inhibitors

Liposome leakage was detected by an increase in fluorescence when Tb^3+^ bound to DPA in Buffer C. Human GSDMD (0.3 µM), dispensed into 384-well plates (Corning 3820) containing PC/PE/CL liposomes (50 µM liposome lipids), was incubated with compounds from the ICCB-Longwood Screening Facility collection for 1 hr before addition of caspase-11 (0.15 µM) to each well. The fluorescence intensity of each well was measured at 545 nm with an excitation of 276 nm 1 hr after addition of caspase-11 using a Perkin Elmer EnVision plate reader. The final percent inhibition was calculated as [(fluorescence_test compound_ − fluorescence_negative control_)/(fluorescence_positive control_ − fluorescence_negative control_)] × 100, where wells with GSDMD without inhibitors was used as positive control and without caspase-11 as negative control. 50% inhibition was arbitrarily chosen as a threshold. The hits were evaluated in concentration-response experiments in a dose range of 0.008–50 µM to determine IC_50_.

### Fluorescent protein labelling and microscale thermophoresis binding assay

His_6_-MBP-GSDMD was labeled with AlexaFluor-488 using the Molecular Probes protein labelling kit. Binding of inhibitors to GSDMD was evaluated using microscale thermophoresis (MST). Ligands (49 nM - 150 μM) were incubated with purified AlexaFluor-488-labeled protein (80 nM) for 30 min in assay buffer (20 mM HEPES, 150 mM NaCl, 0.05% Tween 20). The sample was loaded into NanoTemper Monolith NT.115 glass capillaries and MST carried out using 20% LED power and 40% MST power. *K*_d_ values were calculated using the mass action equation and NanoTemper software.

### Caspase-1 and caspase-11 inhibition assays

The fluorogenic assay for caspase-1 and caspase-11 activity is based on release of 7-amino-4-methylcoumarin (AMC) from the caspase substrate Ac-YVAD-AMC. Compounds (8 nM - 50 μM) were incubated with 0.5 U of caspase-1 or caspase-11 for 30 min in assay buffer (20 mM HEPES, 150 mM NaCl) in 384-well plates (Corning 3820) before addition of Ac-YVAD-AMC (40 μM) to initiate the reactions. Reactions were monitored in a SpectraMax M5 plate reader (Molecular Devices, Sunnyvale, California USA) with excitation/emission wavelengths at 350/460 nm. The fluorescence intensity of each reaction was recorded every 2 min for 2 hrs.

### High-throughput cell viability assay

THP-1 cells seeded at a density of 4000 cells per well in 96-well plates (Corning 3610), were differentiated by exposure to 50 nM PMA for 36 hrs before being primed with 100 ng/mL LPS. Primed THP-1 cells were pretreated with each compound for 1 hr before addition of 20 μM nigericin or medium as control. The number of surviving cells was determined by CellTiter-Glo assay 1.5 hrs later. The final percent cell viability was calculated using the formula [(luminescence_test compound_ − luminescence_negative control_)/(luminescence_positive control_ − luminescence_negative control_)] × 100, where wells with only LPS were used as positive controls and wells treated with LPS and nigericin as negative controls. The IC_50_ of each compound in the cell viability assay was determined by concentration-response experiments in a dose range of 0.39 - 50 μM.

### Mass spectrometry and sample preparation

Gel bands were cut into 1 mm size pieces and placed into separate 1.5 mL polypropylene tubes. 100 μl of 50% acetonitrile in 50 mM ammonium bicarbonate buffer were added to each tube and the samples were then incubated at room temperature for 20 min. This step was repeated if necessary to destain gel. Then, the gel slice was incubated with 55 mM iodoacetamide (in 50 mM ammonium bicarbonate) for 45 min in the dark at room temperature, before the gel was washed sequentially with 50 mM ammonium bicarbonate, water and acetonitrile. Samples were then dried in a Speedvac for 20 min. Trypsin (Promega Corp.) (10 ng/μL in 25 mM ammonium bicarbonate, pH 8.0) was added to each sample tube to just cover the gel, and samples were then incubated at 37 °C for 6 hrs or overnight.

After digestion, samples were acidified with 0.1% formic acid (FA) and 3 μl of tryptic peptide solution was injected. Nano-LC/MS/MS was performed on a Thermo Scientific Orbitrap Fusion system, coupled with a Dionex Ultimat 3000 nano HPLC and auto sampler with 40 well standard trays. Samples were injected onto a trap column (300 μm i.d. × 5mm, C18 PepMap 100) and then onto a C18 reversed-phase nano LC column (Acclaim PepMap 100 75 μm × 25 cm), heated to 50 °C. Flow rate was set to 400 nL/min with 60 min LC gradient, using mobile phases A (99.9% water, 0.1% FA) and B (99.9% acetonitrile, 0.1% FA). Eluted peptides were sprayed through a charged emitter tip (PicoTip Emitter, New Objective, 10 +/- 1 μm) into the mass spectrometer. Parameters were: tip voltage, +2.2 kV; Fourier Transform Mass Spectrometry (FTMS) mode for MS acquisition of precursor ions (resolution 120,000); Ion Trap Mass Spectrometry (ITMS) mode for subsequent MS/MS via higher-energy collisional dissociation (HCD) on top speed in 3 s.

Proteome Discoverer 1.4 was used for protein identification and modification analysis. UniPort human database was used to analyze raw data. Other parameters include the following: selecting the enzyme as trypsin; maximum missed cleavages = 2; dynamic modifications are carbamidomethyl (control), diethyldithiocarbamate (from C-23) and Bay 11-7082 on cysteine; oxidized methionine, deaminated asparagine and glutamine; precursor tolerance set at 10 ppm; MS/MS fragment tolerance set at 0.6 Da; and +2 to +4 charged peptides are considered. Peptide false discovery rate (FDR) was set to be smaller than 1% for significant match.

### Cell lines and treatments

THP-1 cells and HEK293T cells (obtained from ATCC) were grown in RPMI with 10% heat-inactivated fetal bovine serum, supplemented with 100 U/ml penicillin G, 100 μg/ml streptomycin sulfate, 6 mM HEPES, 1.6 mM L-glutamine, and 50 μM 2ME. C57BL/6 mouse iBMDM cells were kindly provided by J. Kagan (Boston Children’s Hospital) and cultured in DMEM with the same supplements. Cells were verified to be free of mycoplasma contamination. Transient transfection of HEK293T cells was performed using Lipofectamine 2000 (Invitrogen) according to the manufacturer’s instructions. iBMDM cells were transfected by nucleofection using the Amaxa Nucleofector kit (VPA-1009). Generally, THP-1 cells were first differentiated by incubation with 50 nM PMA for 36 hrs and then primed with LPS (1 μg/ml) for 4 hrs before treatment with nigericin (20 μM). To examine IκBα phosphorylation and degradation as well as IL-1β induction, PMA-differentiated THP-1 cells were stimulated with LPS (1 μg/ml) for 0.5, 1 and 4 hrs, respectively. For noncanoical inflammasome activation, 1 million iBMDM cells were electroporated with 1μg ultra LPS.

### Cytotoxicity and cell viability assays

Cell death and cell viability were determined by the lactate dehydrogenase release assay using the CytoTox 96 Non-Radioactive Cytotoxicity Assay kit (Promega) and by measuring ATP levels using the CellTiter-Glo Luminescent Cell Viability Assay (Promega), respectively, according to the manufacturer’s instructions. Luminescence and absorbance were measured on a BioTek Synergy 2 plate reader.

### Immunoblot analysis

Cell extracts were prepared using RIPA buffer (50 mM Tris-HCl pH 7.4, 150 mM NaCl, 1 mM EDTA, 1% Triton X-100, 0.1% SDS, 0.5% deoxycholate) supplemented with a complete protease inhibitor cocktail (Roche) and a PhosSTOP phosphatase inhibitor cocktail (Roche). Samples were subjected to SDS-PAGE and the resolved proteins were then transferred to a PVDF membrane (Millipore). Immunoblots were probed with indicated antibodies and visualized using a SuperSignal West Pico chemiluminescence ECL kit (Pierce).

### Caspase-1 activity assay in cells

To measure caspase-1 activation, THP-1 cells were seeded into 96-well plates and differentiated with PMA. After the indicated treatments, cells were incubated with a fluorescent active caspase-1 substrate FAM-YVAD-FMK (Immunochemistry Technologies). Samples were read on a BioTek Synergy 2 plate reader.

### Measurement of cytokines

Concentrations of IL-1β in culture supernatants or mouse serum were measured by ELISA kit (R&D Systems) according to the manufacturer’s instructions.

### Immunostaining and confocal microscopy

Cells grown on coverslips were fixed for 15 min with 4% paraformaldehyde in PBS, permeabilized for 5 min in 0.1% Triton X-100 in PBS and blocked using 5% BSA for 1 hr. Then, cells were stained with the indicated primary antibodies followed by incubation with fluorescent-conjugated secondary antibodies (Jackson ImmunoResearch). Nuclei were counterstained with DAPI (4,6-diamidino-2-phenylindole) (Sigma-Aldrich). Slides were mounted using Aqua-Poly/Mount (Dako). Images were captured using a laser scanning confocal microscope (Olympus Fluoview FV1000 Confocal System) with a 63× water immersion objective and Olympus Fluoview software (Olympus). All confocal images are representative of three independent experiments.

### Statistics

Student’s t-test was used for the statistical analysis of two independent treatments. Mouse survival curves and statistics were analyzed using the Mantel-Cox Log-rank test.

---

**Extended Data Figure 1.**
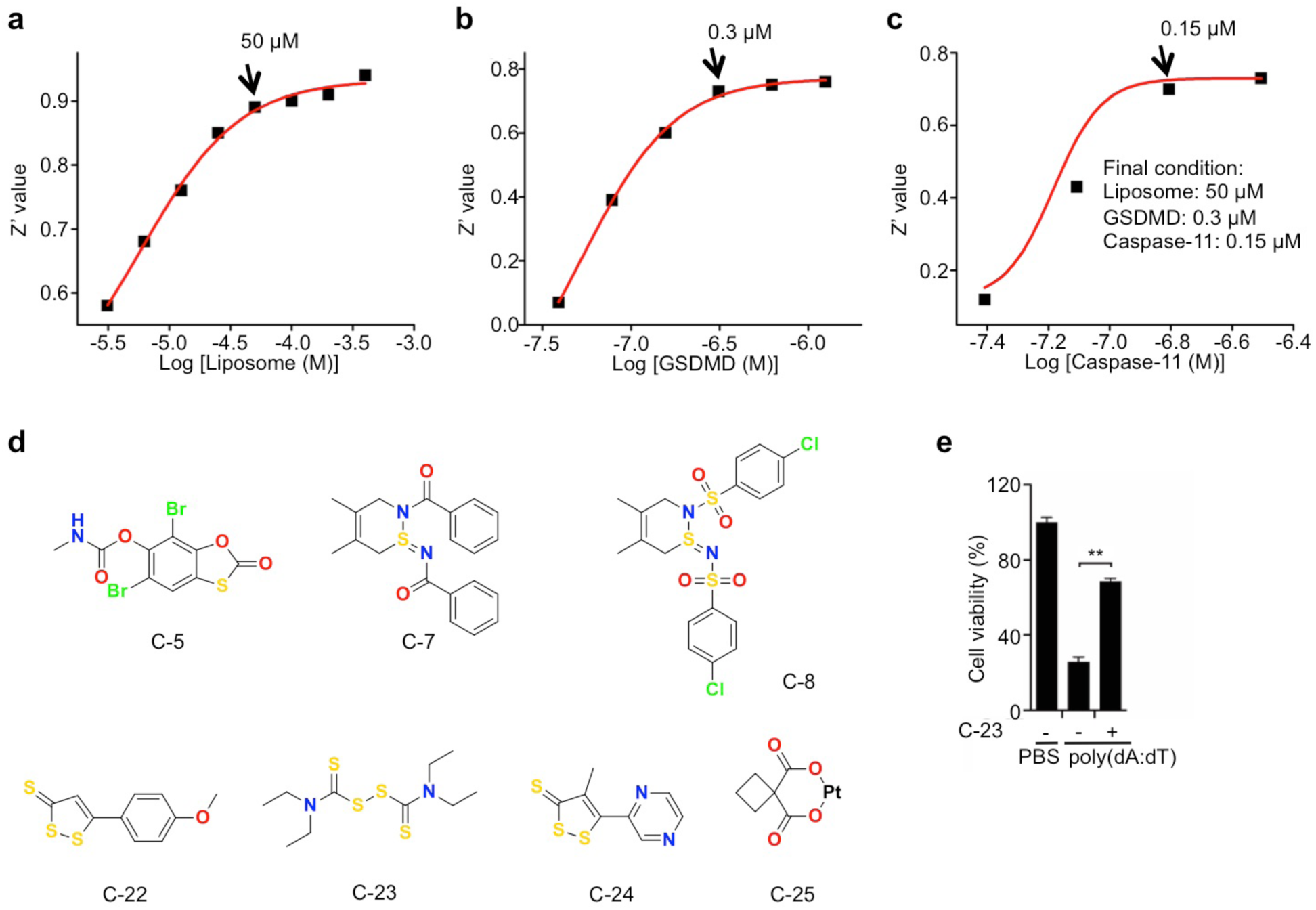
Optimization of the liposome leakage assay. **a**, GSDMD (2.5 μM) and caspase-11 (2.5 μM) were incubated in liposome solutions at various concentrations in 20 mM HEPES buffer (150 mM NaCl) for 1 hr. The concentration of liposome lipids for the screen was set at 50 μM. **b**, Different concentrations of GSDMD and caspase-11 (1:1 ratio) were incubated in liposome (50 μM) solutions for 1 hr. The concentration of GSDMD used in the screen was set at 0.3 μM. **c**, Different concentrations of caspase-11 and GSDMD (0.3 μM) were incubated in liposome (50 μM) solutions for 1 hr. The concentration of caspase-11 used in the screen was set at 0.15 μM. The fluorescence intensity at 545 nm was measured after excitation at 276 nm. **d**, Hit compounds evaluated in binding and/or cell-based assays. **e**, Mouse iBMDMs were pretreated or not with 30 μM disulfiram (C-23) for 1 hr before transfection with PBS or poly(dA:dT) and analysed for cell viability by CellTiter-Glo assay 4 hrs later. Graph shows mean ± s.d; data are representative of three independent experiments. **P < 0.01.

**Extended Data Figure 2.**
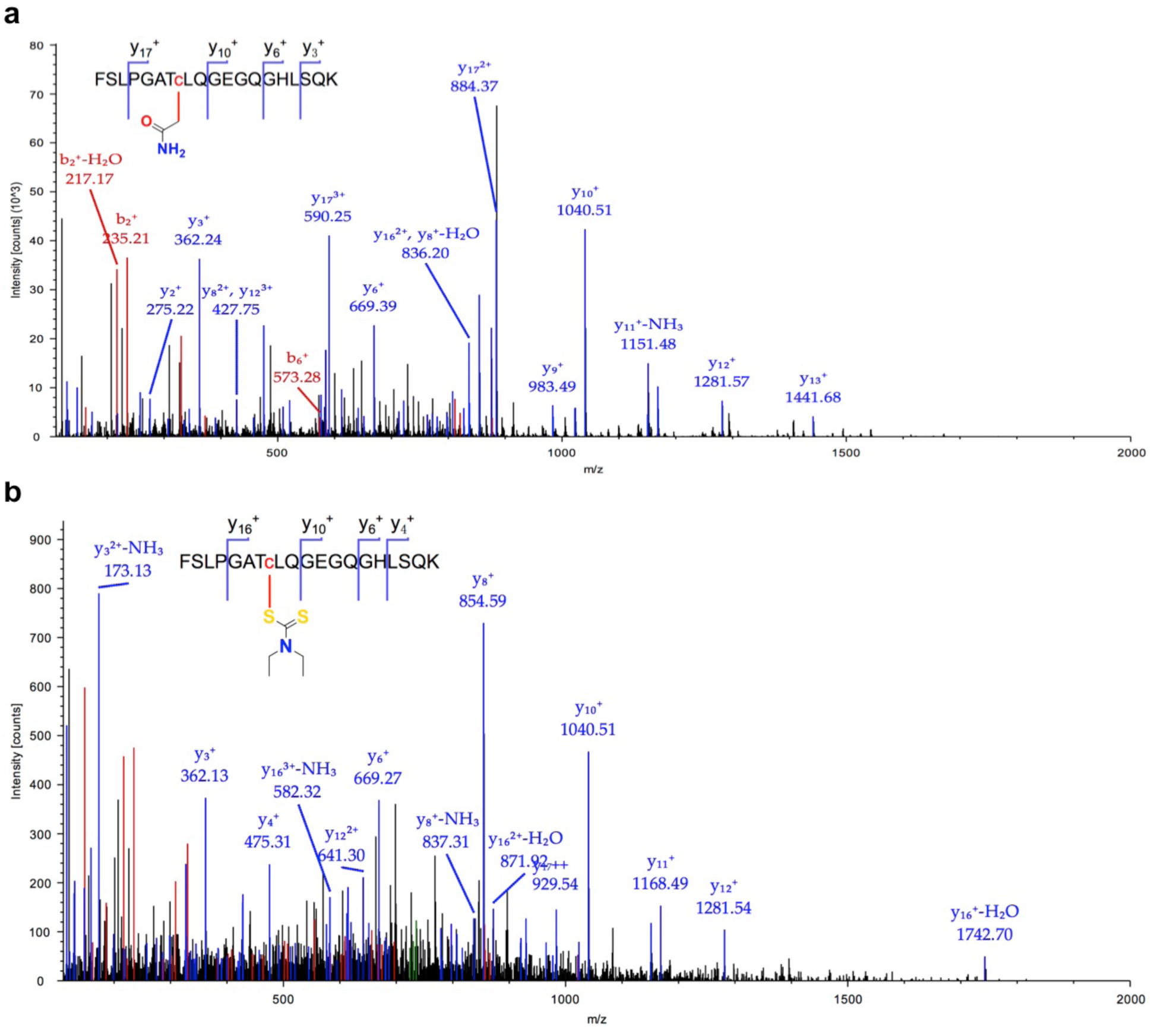
MS/MS spectrum for the peptide containing Cys191 in human GSDMD. **a**, MS/MS spectrum for peptide FSLPGAT**C**LQGEGQGHLSQK modified on cysteine (red) by carbamidomethyl. Protein coverage is 73%. **b**, MS/MS spectrum for peptide FSLPGAT**C**LQGEGQGHLSQK modified on cysteine (red) by C-23. Protein coverage is 72%.

**Extended Data Figure 3.**
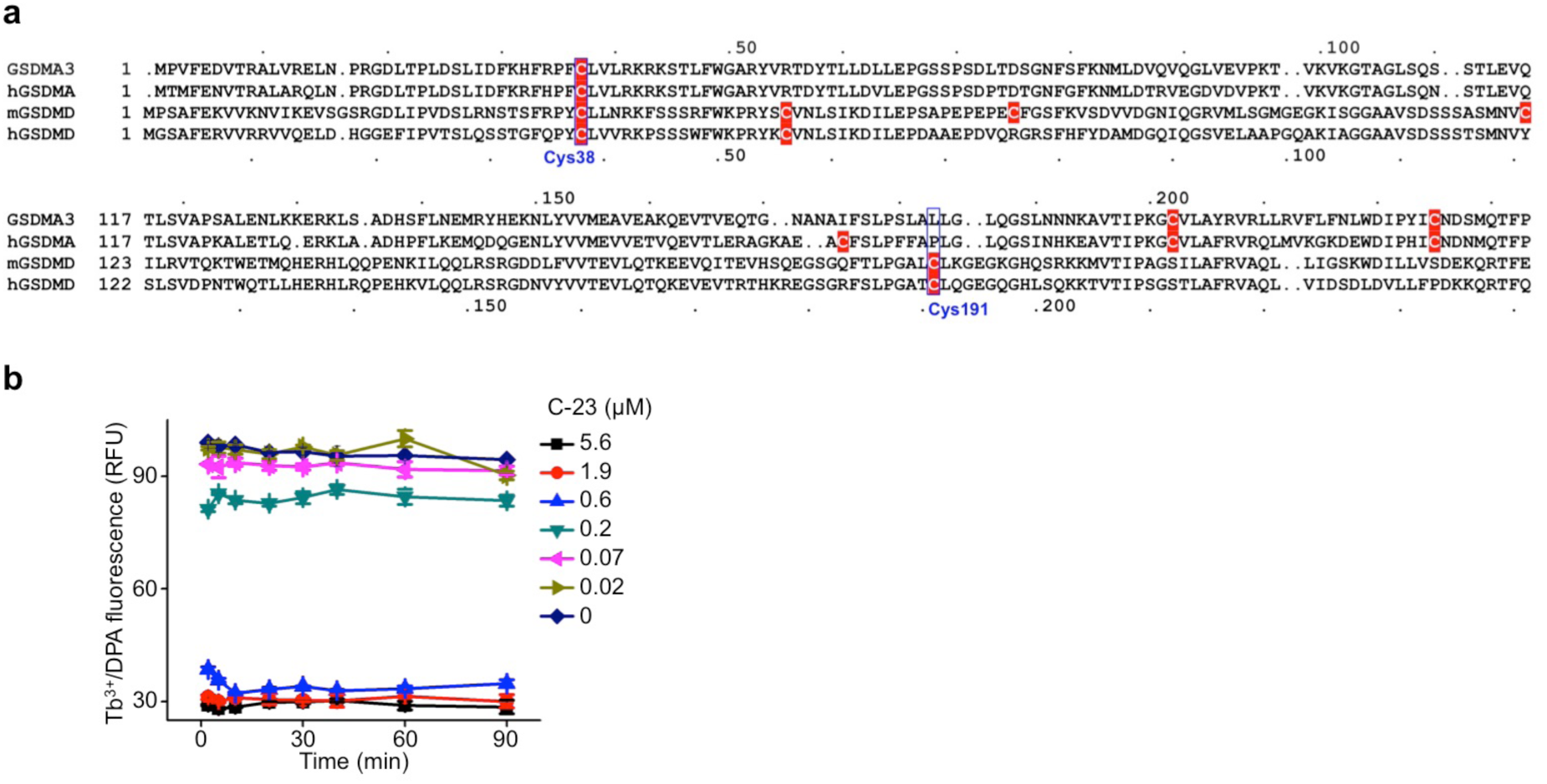
Disulfiram covalently modifies GSDMD Cys191. **a,** Sequence alignment of mouse GSDMA3, human GSDMA (hGSDMA), mouse GSDMD (mGSDMD) and human GSDMD (hGSDMD) showing Cys residues (highlighted in red)**. b**, GSDMD (0.3 μM) was preincubated with the indicated concentrations of C-23 (0 – 5.6 μM) for different durations (2-90 min) before caspase-11 (0.15 μM) in liposome (50 μM) was added.

**Extended Data Figure 4.**
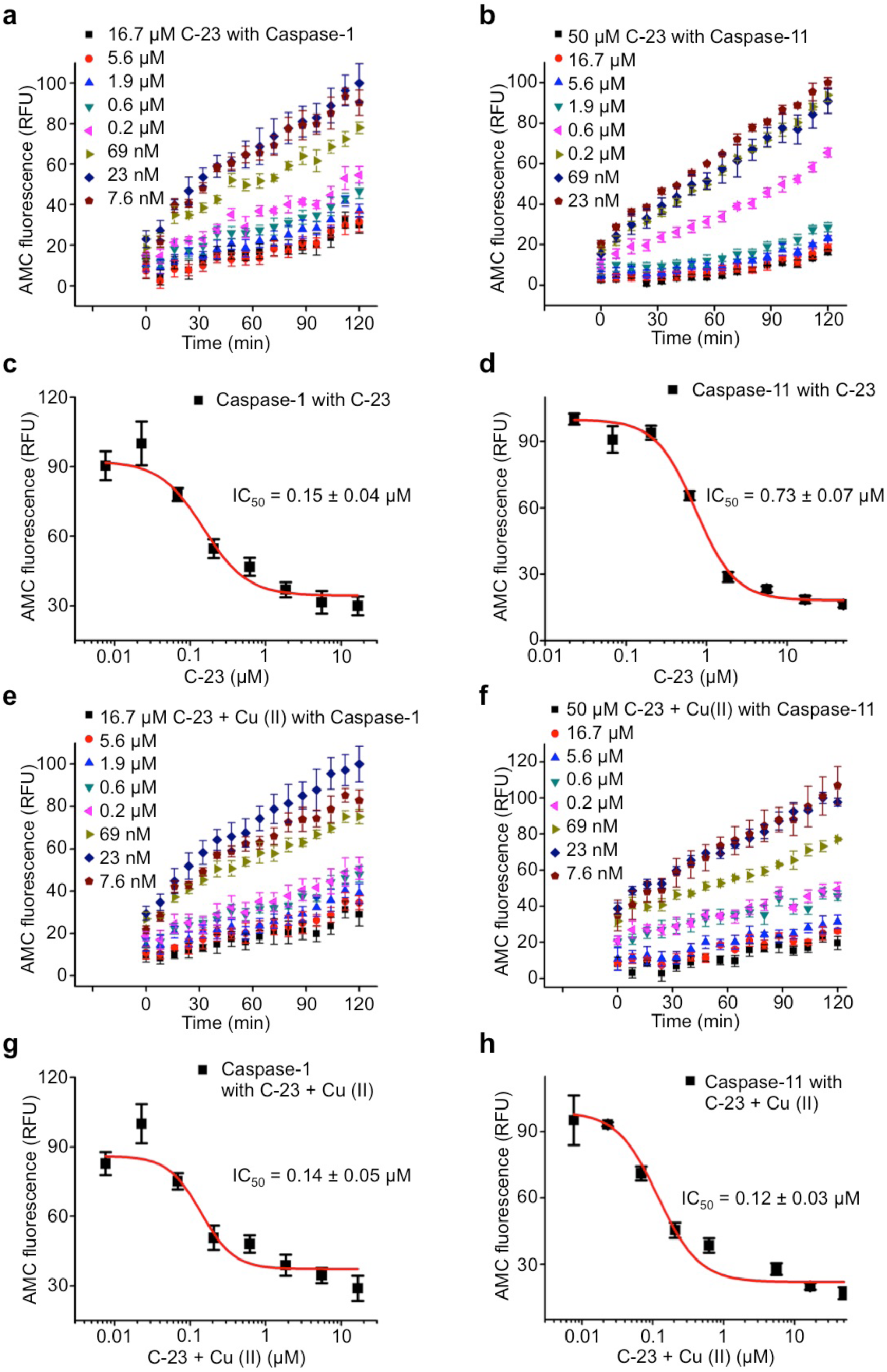
Disulfiram (C-23) inhibits caspase-1 and caspase-11. **a**,**b**, Time course of caspase-1 (**a**) and caspase-11 (**b**) activity in the presence of indicated concentrations of compound C-23. Caspases (0.5 U) were incubated with compound C-23 (at indicated concentrations for 1 hr before adding Ac-YVAD-AMC (40 μM)). **c**,**d**, Dose response curve of compound C-23 in the caspase-1 (**c**) and caspase-11 (**d**) activity assay. **e**,**f**, Time course of caspase-1 (**e**) and caspase-11 (**f**) activity in the presence of indicated concentrations of compound C-23 + Cu(II). Caspases (0.5 U) were incubated with compound C-23 + Cu(II) (at indicated concentrations for 1 hr before adding Ac-YVAD-AMC (40 μM)). **g**,**h**, Dose response curve of compound C-23 in the caspase-1 (**g**) and caspase-11 (**h**) activity assay. Fluorescence intensity at 460 nm was measured after excitation at 350 nm.

**Extended Data Figure 5.**
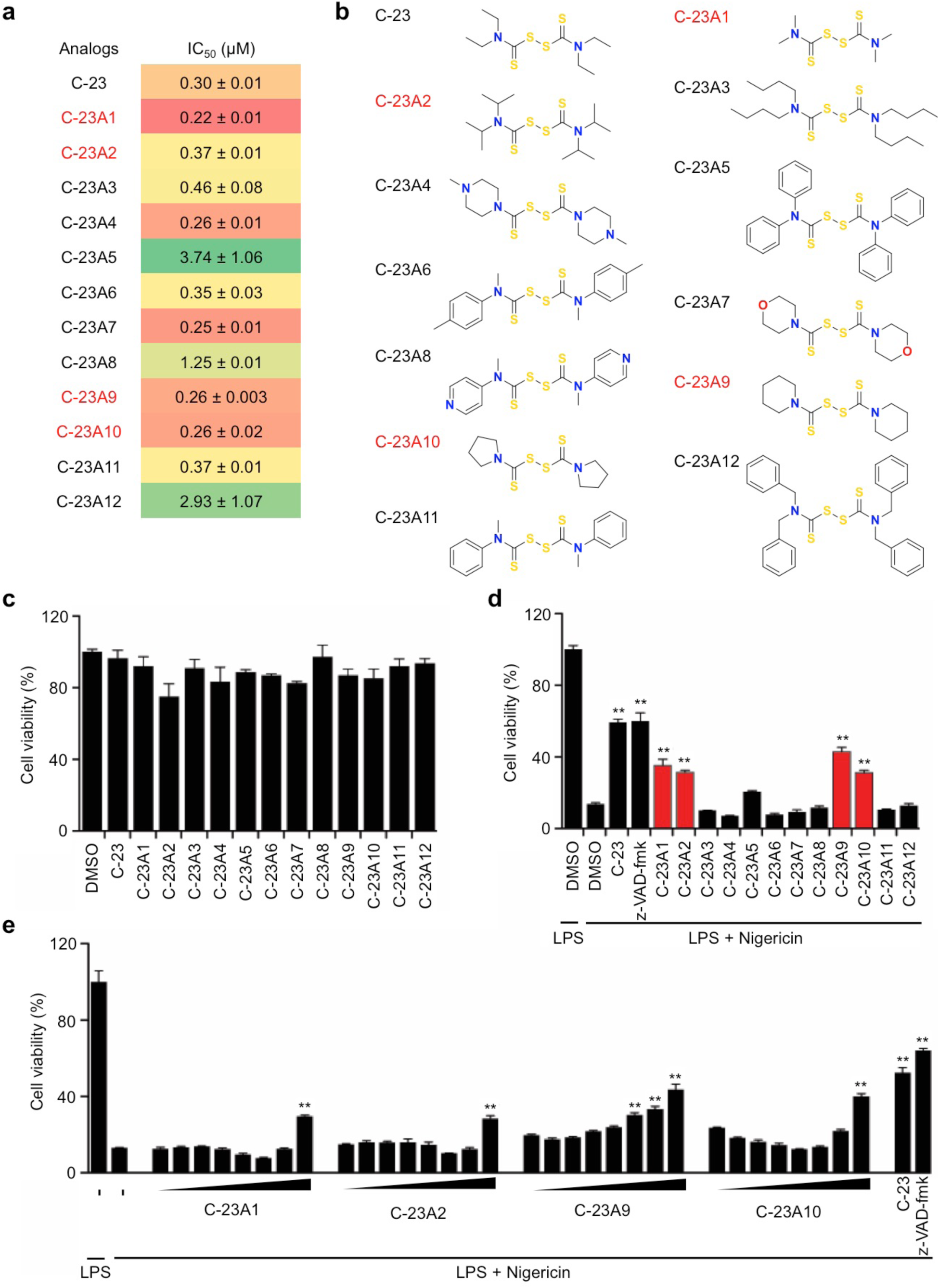
Some disulfiram analogues inhibit GSDMD. **a**, Summary of IC50 of tested disulfiram analogues in the liposome leakage assay. **b**, Structures of analogues. **c**, PMA-differentiated LPS-primed THP-1 cells were treated with the indicated compounds (40 μM) for 3 hrs and tested for viability by CellTiter-Glo assay. **d**, PMA-differentiated LPS-primed THP-1 cells, pretreated with 40 μM disulfiram or the indicated analogues or z-VAD-fmk for 1 hr before treatment or not with nigericin, were assessed for cell viability by CellTiter-Glo assay 2 hrs after adding nigericin. **e**, PMA-differentiated LPS-primed THP-1 cells, pretreated with 40 μM disulfiram or z-VAD-fmk or with 2-fold serial dilutions (concentration range, 0.39-50 μM) of indicated analogues for 1 hr before adding nigericin, were assessed for cell viability by CellTiter-Glo assay 2 hrs after adding nigericin. Graphs show the mean ± s.d. and data shown are representative of three independent experiments. **P < 0.01.

**Extended Data Figure 6.**
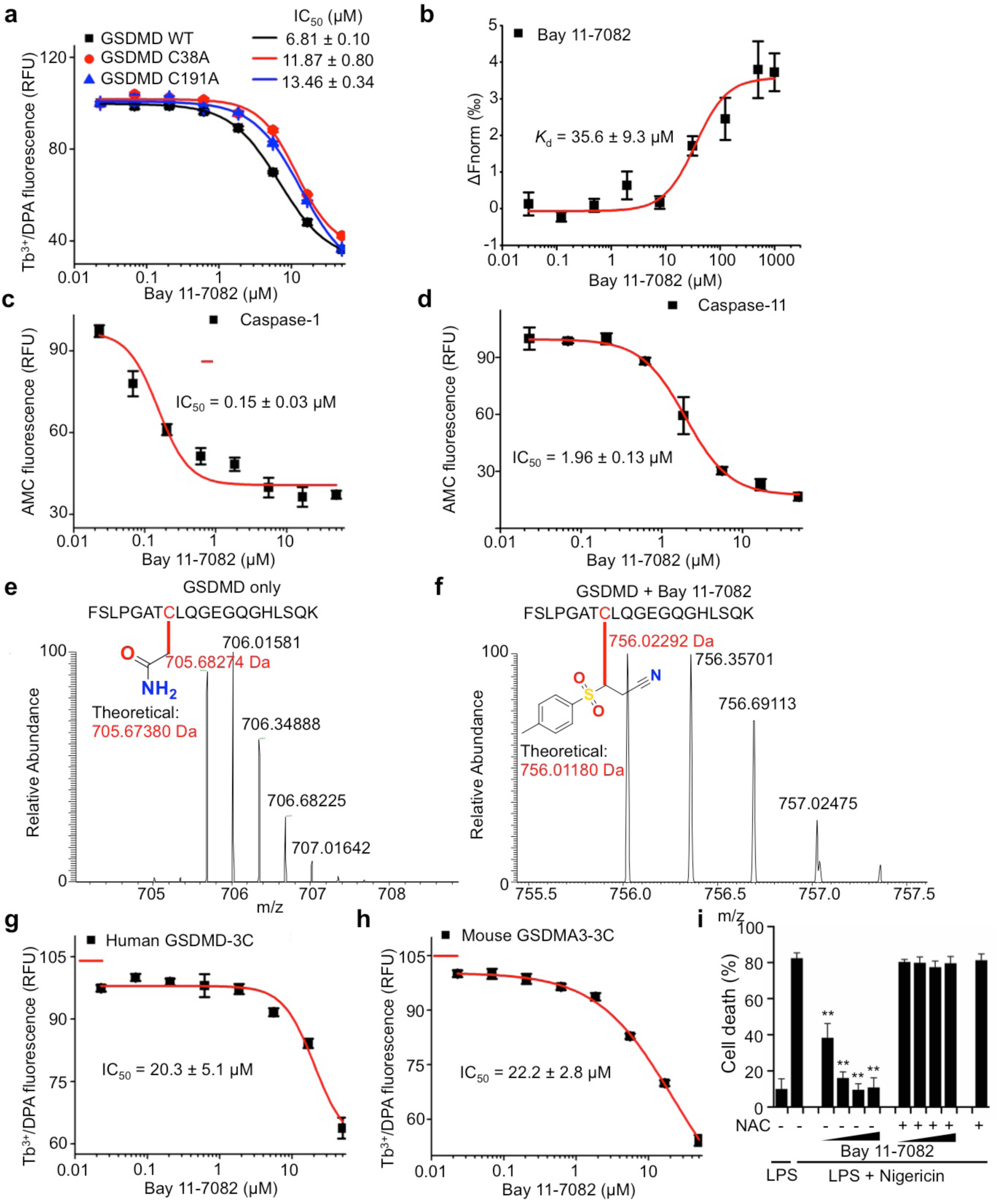
Bay 11-7082 inhibits GSDMD, caspase-1 and caspase-11. **a**, Bay 11-7082 dose response curve of inhibition of liposome leakage by wild-type, C38A or C191A GSDMD (0.3 μM) plus caspase-11 (0.15 μM). **b**, MST measurement of the direct binding of Alexa 488-labeled His-MBP-GSDMD (80 nM) with Bay 11-7082 by NanoTemper. **c**,**d**, Dose response curve of the effect of Bay 11-7082 on caspase-1 (**c**) and caspase-11 (**d**) activity against a fluorescent peptide substrate. **e**,**f**, MS/MS spectra of the Cys191-containing GSDMD peptide FSLPGAT**C**LQGEGQGHLSQK (aa 184-103; 2057.00 Da) modified on Cys191 (red) by carbamidomethyl (an increase of 57.0214 Da) [LC retention time, 22.85 min; a triplet charged precursor ion m/z 705.6827 (mass: 2114.0481 Da; delta M 2.27 ppm) was observed] (**e**) or of the corresponding GSDMD peptide after GSDMD incubation with Bay 11-7082, which was modified on Cys191 (red) (an increase of 207.0354 Da). [LC retention time, 17.20 min; a triplet charged precursor ion m/z 756.0229 (mass: 2264.0688 Da; delta M 11.7 ppm) was observed.] (**f)**. **g**,**h**, Dose response curve of the effect of Bay 11-7082 on liposome leakage induced by 0.3 μM human GSDMD-3C (**g**) or mouse GSDMA3-3C (**h**) plus 0.15 μM 3C protease. **i**, Effect of 1 hr preincubation of Bay 11-7082 with N-acetylcysteine (NAC, 500 μM) on inhibition of pyroptosis of LPS + nigericin treated THP-1 cells. 2-fold dilutions of Bay 11-7082 from 5-40 μM were used. Graphs show the mean ± s.d. and data shown are representative of three independent experiments. **P < 0.01.

**Extended Data Figure 7.**
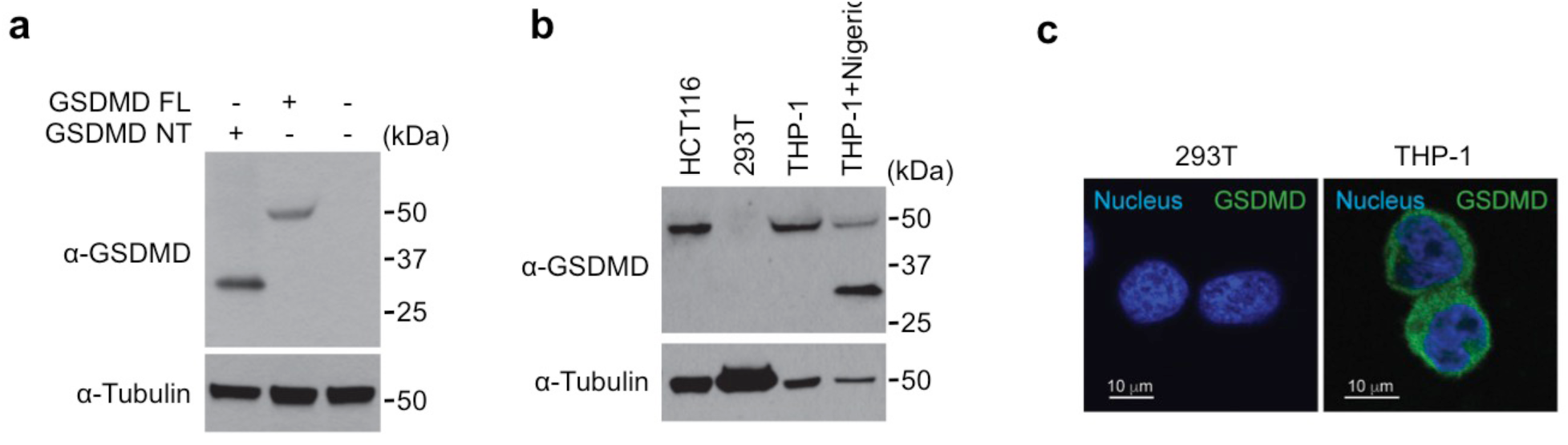
Mouse monoclonal antibody recognizes full-length human GSDMD and the GSDMD-NT pore form on immunoblots and by immunofluorescence microscopy. The monoclonal antibody against GSDMD was generated by immunizing mice with recombinant human GSDMD and boosting with recombinant human GSDMD-NT as described in **Methods**. **a,** HEK293T cells were transfected with the indicated plasmids and cell lysates were analysed by immunoblot of reducing gels probed with the indicated antibodies. **b,** Cell lysates of HCT116, 293T and THP-1 cells, treated or not with nigericin, were immunoblotted with the indicated antibodies. 293T cells do not express endogenous GSDMD. **c,** 293T and THP-1 cells were immunostained with the anti-GSDMD monoclonal antibody and co-stained with DAPI (blue). 293T cells that do not express GSDMD show no background staining.

